# Causal SNP regulating *FAM13B* expression identified for the Chr. 5q31 atrial fibrillation susceptibility locus

**DOI:** 10.1101/719914

**Authors:** Gregory Tchou, Shamone Gore-Panter, Jeffrey Hsu, Fang Liu, Nana Liu, Christine S. Moravec, John Barnard, David R. Van Wagoner, Mina K. Chung, Jonathan D. Smith

**Author notes:** These authors are all corresponding authors Address correspondence to Jonathan D. Smith, Box NC10, Cleveland Clinic, 9500 Euclid Avenue, Cleveland, OH, 44195, USA. These authors contributed equally to this study (Gregory Tchou, Shamone Gore-Panter, Jeffrey Hsu, Fang Liu). **Abbreviations:** GWAS, genome wide association study; AF, atrial fibrillation; SNP, single nucleotide polymorphism; eQTL, expression quantitative trait loci; LA, left atrium; RNAseq, next generation RNA sequencing; lncRNAs, long noncoding RNA; FDR, false discovery rate; AFR, atrial fibrillation rhythm at time of surgery; SR, sinus rhythm at time of surgery; SVA, surrogate variable analysis; VST, variance-stabilized transformation; MDS, multidimensional scaling; LAA, left atrial appendage; AF/SR, history of AF but in sinus rhythm at time of surgery; TSS, transcription start site; LD, linkage disequilibrium; iPSC, inducible pluripotent stem cell; I_Na_, sodium current; I_NaL_, late sodium current.

## Abstract

**Rationale:** Our prior RNA sequencing study found that FAM13B gene expression in human left atrial appendages was strongly associated with an atrial fibrillation (AF) susceptibility-associated variant on chr. 5q31.

**Objective:** To identify the common genetic variant responsible for regulating FAM13B expression and the effect of FAM13B expression on cardiomyocyte gene expression in order to gain insight into the functional mechanism of the chr. 5q31 AF susceptibility locus.

**Methods and Results:** By taking advantage of a smaller linkage disequilibrium block in African descent subjects and available chromatin conformation data, we identified the common single nucleotide polymorphism (SNP) rs17171731 as a candidate genetic variant controlling FAM13B gene expression in the left atrium. Functional analysis demonstrated that the AF risk allele of rs17171731 had less enhancer activity than the protective allele. Gel mobility shift studies determined that the risk allele bound to an additional protein that may function as a transcriptional repressor. Knockdown of *FAM13B* expression in stem cell-derived human cardiomyocytes (iCM) altered the expression of >1000 genes and modified the sodium current, consistent with increased susceptibility to atrial fibrillation. Transfection of GFP tagged FAM13B into iCMs demonstrated expression on the plasma membrane and at the Z-disk.

**Conclusions:** The chr. 5q31 AF risk variant was identified as rs17171731, with the risk allele having less enhancer activity, leading to decreased expression of *FAM13B*, which resides on the plasma membrane and the Z-disk, and appears to play a role in the regulation of cardiomyocyte gene expression and the late sodium current.

---

Atrial fibrillation (AF) is the most common human arrhythmia and has a strong heritable component. ^1^ The most recent genome wide association studies (GWAS) for AF susceptibility identified ∼100 loci associated with AF. ^2,3^. However, further work on each locus is necessary to identify the causal gene, the causal genetic variant, and the mechanism for association with AF. Since most AF GWAS SNPs, or those in strong linkage disequilibrium with them, do not alter the protein amino acid sequence, we hypothesize that most of these GWAS loci are regulatory, leading to changes in gene expression rather than protein structure. Our prior expression quantitative trait locus (eQTL) study utilized next generation RNA sequencing (RNAseq) and single nucleotide polymorphism (SNP) genotyping in human left atrial appendages to identify the *cis* genes whose expression are associated with the top SNP at the AF GWAS loci. ^4^ We identified FAM13B as the gene whose expression is associated with the top GWAS SNP at chr. 5q31. Here, we describe how an eQTL analysis in a small African descent cohort led to the identification of a candidate SNP, rs17171731, in the chr. 5q31 AF GWAS locus that controls *FAM13B* gene expression. We validated this candidate SNP through reporter gene transfections, gel mobility shift assays, and gene editing in human pluripotent stem cells. Thus, the chr. 5q31 GWAS locus appears to be mediated by regulating the expression of FAM13B, a poorly characterized member of the rho GTP activating domain gene family. ^5,6^ We then used siRNA to knockdown *FAM13B* in stem cell-derived cardiomyocytes in order to investigate the mechanism by which decreased *FAM13B* gene expression leads to increased AF risk. Knockdown of *FAM13B* expression altered the expression of >1000 cardiomyocyte genes, including a subunit of the voltage-dependent sodium channel, which suggests possible mechanisms by which the genetic regulation of *FAM13B* could result in AF susceptibility. Expression of GFP-tagged FAM13B in stem cell derived cardiomyocytes s localized FAM13B protein to the plasma membrane and Z-disk.

## Methods

### Human Left Atrial Tissue Samples and Processing

Human left atrial appendage tissues were obtained from patients undergoing elective surgery to treat AF, valve disease, or other cardiac disorders. LA tissue specimens were also obtained from non-failing donor hearts not used for transplant. Demographics of this population were previously published. ^4^ Total RNA was isolated for subsequent sequencing analysis. These methods are detailed in our prior publication. ^4^

### Genomic DNA isolation and SNP microarray

As previously described ^4^, genomic DNA was genotyped using Illumina Hap550v3 and Hap610-quad SNP microarrays. In order to derive genotypes in addition to those on the microarray, the SNP data was imputed to 1000 Genomes Project phase 2 to generate genotypes for ∼19 million SNPs, using the IMPUTE software. ^7^

### RNA isolation, sequencing, and analysis

As previously described ^4^, 100-bp paired-end sequencing was performed on the Illumina HiSeq 2000 platform and multiplexed to 6 samples across two lanes. The sequence reads were mapped to the human genome to derive a digital count of the expression of genes, which were defined using the Ensembl gene catalog (version 71). Reads were quantile normalized, and gene counts for eQTL analysis were variance-stabilized transformed as previously described.^4^ eQTL analyses were performed separately for each racial group with beta-coefficients calculated as the additive effect of one allelic difference on log_2_ gene expression. Genetic multidimensional scaling (MDS) from SNP array genotyping were additionally calculated and added as covariates for the eQTL calculation. matrixeQTL ^8^ was used to test associations between genotype and variance stabilized counts. The qvalue package was used to calculate false discovery rate (FDR) from the complete list of p values. ^9^

### Human stem cell culture and cardiac differentiation

H9 human embryonic stem cells (WiCell) were cultured in TesR-E8 media (STEMCELL 05990) on plates coated with growth factor-reduced Matrigel (ThermoFisher CB-40230). After passaging, H9 cells were cultured in the presence of 10 μM Y-27632 (Abcam) for 24-48 hours. Cardiac differentiation of H9 cells was conducted using the STEMdiff Cardiomyocyte Differentiation Kit (STEMCEll 05010).

### Reporter gene expression analysis

26-mer oligonucleotides (Table 1) containing the *FAM13B* SNP rs17171731 reference or risk allele sequences were cloned into the luciferase reporter pT81luc. ^10^ The luciferase reporter and the β-galactosidase expression vector pCH110 were transfected into H9 human embryonic stem cell-derived cardiomyocytes via electroporation (Nucleofector-II program A-23, Lonza VPH-5012). Cell lysates were prepared and analyzed for β-galactosidase and luciferase activities using the Dual Light Reporter Gene Assay system (ThermoFisher T1003).

**Table 1.**
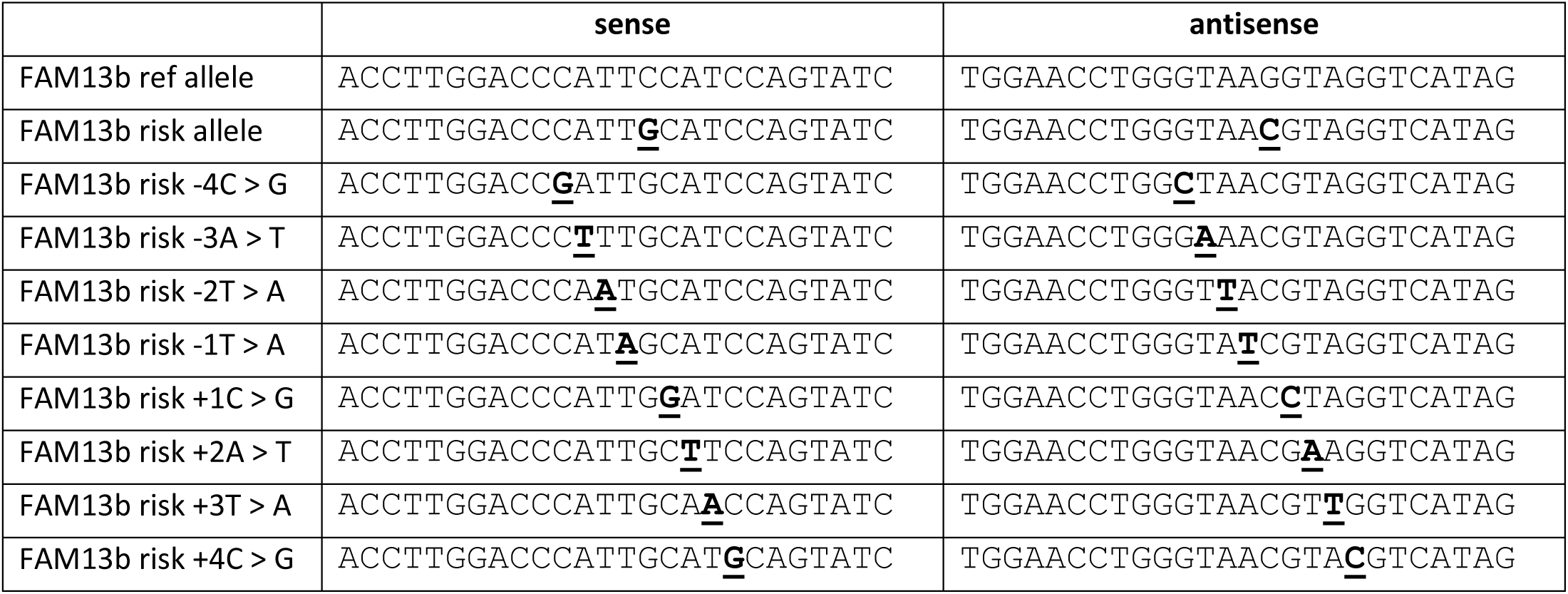
Oligonucleotides used for FAM13B enhancer screen and gel mobility shift assays.

### FAM13B electrophoretic mobility shift assays

26-mer oligonucleotides containing the rs17171731 reference or risk allele sequences (Table 1) were labeled with γ-^32^P-ATP (10 μCi/μL) using T4 polynucleotide kinase (NEB M0201) and annealed with complementary oligonucleotides in 10 mM Tris, 1 mM EDTA, 50 mM NaCl. The resulting probes were combined on ice with human heart nuclear extract (Active Motif 36041), 1.25 μg poly(dI dC) (Sigma-Aldrich P4929) and reaction buffer (final concentration 45 mM KCl, 1 mM MgCl_2_, 15 mM HEPES pH 7.9, 1 mM DTT, 3% Ficoll 400) in the presence or absence of unlabeled competitor oligonucleotides, then incubated at room temperature for 25 minutes before running on Novex 6% DNA Retardation gels (ThermoFisher EC63655BOX). Dried Gels were exposed to X-ray film. We attempted to identify proteins bound to the probes using the following antibodies: ABI3, ATF4, MTF-1, POU2F1, POU2F2, SOX4, and SOX8 (One World Lab), as well as ZNF143 (R&D Systems).

### IPSC-derived cardiomyocyte culture and siRNA knockdown of *FAM13B*

Cardiomyocytes derived from human inducible pluripotent stem cells (iPSCs) (iCell cardiomyocytes, Cellular Dynamics International, Madison, WI) were cultured and siRNA knockdown was performed using Cellular Dynamics International protocols. In short, 1.5 × 10^5^ cells were plated per well in fibronectin (Sigma)-coated 12-well plates. The cells were cultured in iCell cardiomyocyte maintenance media (Cellular Dynamics Int.) for seven days; beating was observed at about day six. Final concentrations of 50 nM/well of control scramble siRNA (Thermo-Fisher/Ambion silencer select cat#4390843) or *FAM13B* siRNA (Thermo-Fisher/Ambion cat# 439242, id S27906) were transfected into iCell cardiomyocytes using the TransIT-TKO transfection reagent (Mirus Bio). The cells were incubated with the siRNA-reagent complexes for 48-96 hours followed by either cellular electrophysiology studies or total RNA isolation, cDNA preparation, qRT-PCR, and RNAseq (described above).

### Quantitative reverse transcriptase-polymerase chain reaction (qRT-PCR)

Total RNA was isolated from the iCell cardiomyocytes after siRNA knockdown using the Cellular Dynamics International protocol (Total RNA protocol) and the RNeasy Micro Kit (Qiagen) with the following modification: from step 3 through 5, the reagent volumes were doubled to accommodate the increased well size. 100-250 μg of RNA was reverse transcribed using SuperScript IV VILO™ Master Mix (Thermo-Fisher/Invitrogen). 12.5 μl of the TaqMan^®^ gene expression master mix (Applied Biosystems), 1.25 μl of FAM-labeled FAM13B primer/probe mix (assay number HS00991421_M1, Thermo-Fisher), and 1.25 μl VIC-labeled primer limited cardiac actin (ACTC1) primer/probe mix (assay number Hs00606316_m1, Thermo-Fisher) were mixed to create a master mix to be added to each sample. 15 μl aliquots of master mix were dispensed into individual wells of a 96-well working plate along with 10 μl of undiluted cDNA. A 5 μl aliquot of this working mixture was pipetted in triplicate to a 384-well assay plate using an epMotion (Eppendorf model 5070) automated pipetting robot. Real time PCR was performed using a Bio-RAD CRX thermocycler that is calibrated for FAM and VIC fluorescent probes. Thermal cycling was performed with a hot-start at 95°C for 10 minutes, followed by 40 cycles of 95°C for 15 seconds and 60°C for 60 seconds. The ΔC(t) values for FAM13B expression levels were calculated relative to ACTC1. ΔΔC(t) values were used to calculate the decrease of FAM13B expression and the results were converted to base 10 values by calculating 2^−ΔΔC(t)^.

### Cellular electrophysiology methods

Patch clamp studies of FAM13B knockdown cells focused on measurement of sodium currents (INa) in scramble and FAM13B-siRNA treated iCell cardiomyocytes. Cells were replated at low density 24 hours after siRNA treatment (non-touching), and electrophysiology studies were performed 48 to 96 hours after transfection. Sodium currents were recorded at room temperature using whole cell clamp techniques. Pipettes (Corning 8161 thin-walled glass, 1.5-3 MΩ) were filled with a solution containing 135 mM CsCl2, 10 mM NaCl, 2 mM CaCl_2_, 5 mM EGTA, 10 mM HEPES, 5 mM MgATP, and titrated to pH 7.2 with CsOH. The cells were continuously superfused with an extracellular solution containing: 50 mM NaCl, 1.8 mM CaCl_2_, 1 mM MgCl_2_, 110 mM CsCl, 10 mM glucose, 10 mM HEPES, 1 μM nifedipine (added fresh daily), pH 7.4. Currents were recorded with an Axopatch 1C amplifier controlled with pClamp 8.1 software (Molecular Devices). After seal formation and patch rupture, whole cell access resistance was 2-5 MΩ. To maintain cell viability, resting potential was held at −60 mV, with more negative steps given 100 ms before depolarizing steps used to record sodium currents. Voltage clamp protocols are shown in Figure 4 A,C,F. Current amplitudes recorded with the steady-state inactivation protocol (Figures 4C,D) were fit with a Boltzmann curve in Clampfit 8.1 (pClamp, Molecular Devices). To avoid time-dependent changes in siRNA exposure, data were obtained from similar numbers of scramble and siRNA-treated cells on each day. Data are reported as mean ± standard error of the mean. All electrophysiologic comparisons between scramble and FAM13B-siRNA treated cells were evaluated using a 2 tailed T-test or a 2-way ANOVA as indicated.

**Figure 1.**
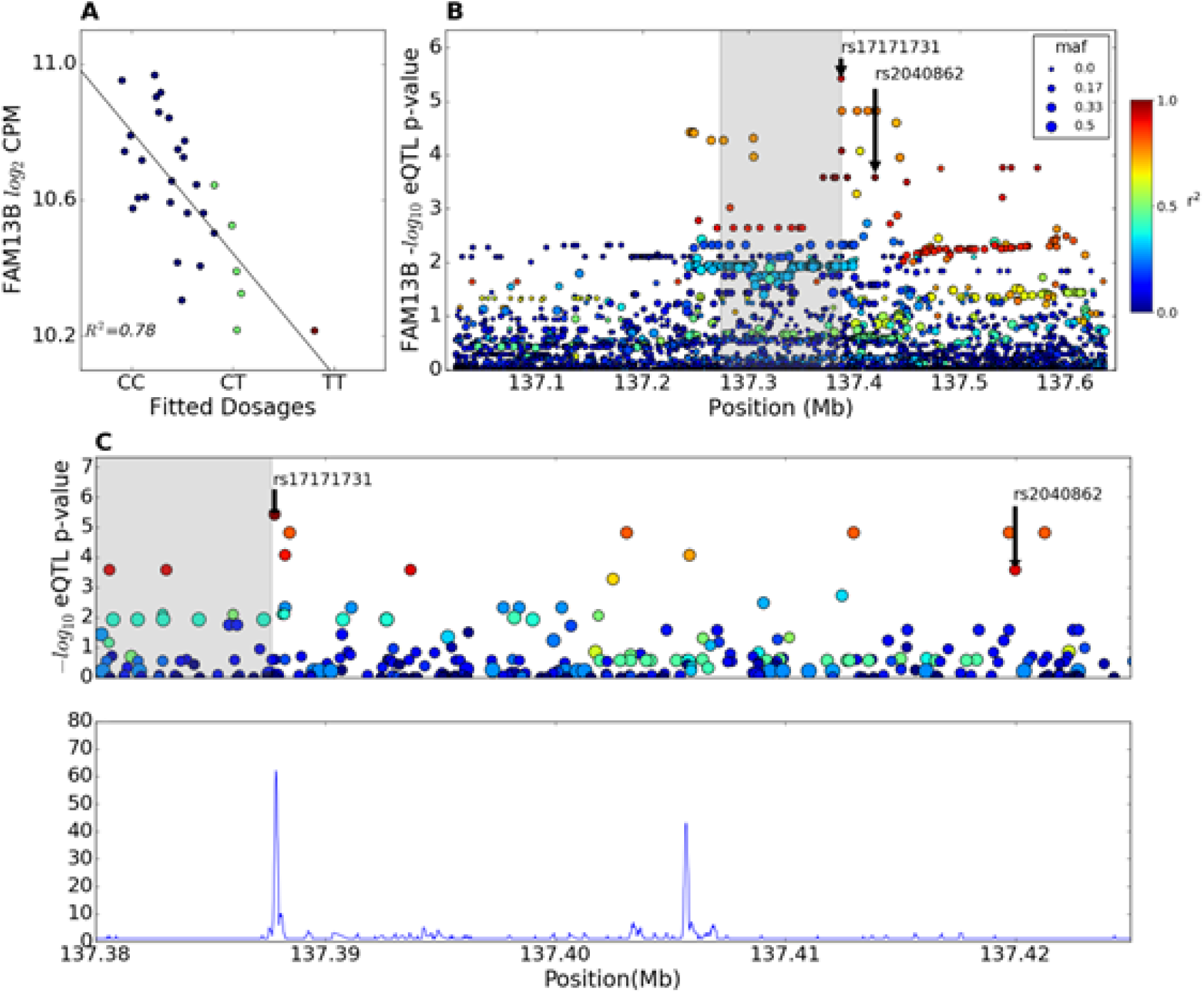
*FAM13B cis*-eQTL. **A.** Partial regression plot showing the effect of rs17171731 on LA expression of the *FAM13B* gene in African American subjects. Blue dots are subjects homozygous for the major allele, green dots are heterozygotes, and red dots are homozygous for the minor allele (r^2^ = 0.78, q-value =0.01). **B.** eQTL p-values and LD relationship with rs17171731 in the region around *FAM13B* in African American subjects (p=2.63×10^-5^). The grey shaded area represents the longest FAM13B gene (transcribed from left to right) in Ensembl, but the LAA transcript starts ∼20 kb proximal to rs17171731. **C.** Zoom in on FAM13B cis-eQTL for African American subjects (top). The SNP rs17171731 occurs in a DNAse hypersensitivity peak while the AF GWAS SNP rs2040862 does not (bottom, Roadmap Epigenomics Project data for human fetal heart).

**Figure 2.**
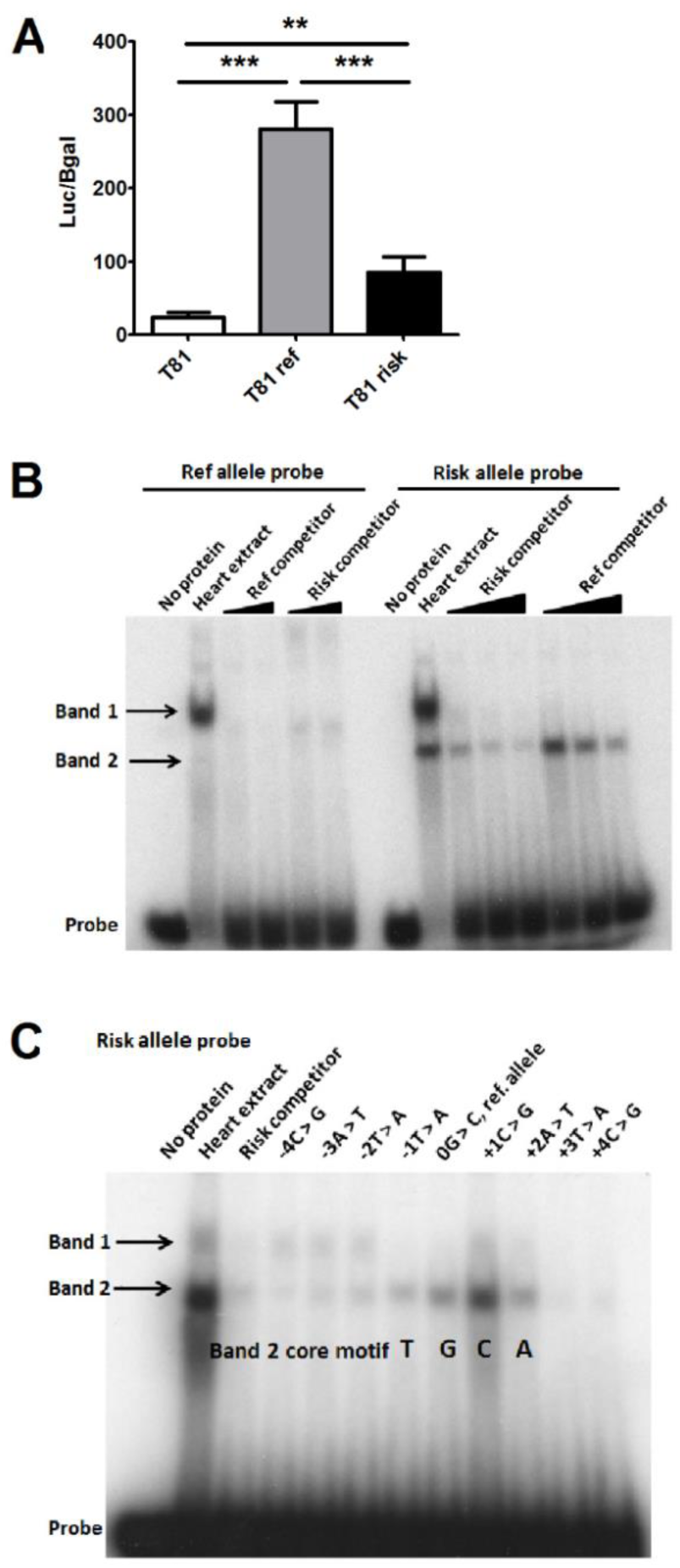
Functional studies for rs17171731, the candidate AF-causal SNP at chr 5q31. **A.** The effects of the reference and AF-risk allele of rs17171731 on luciferase expression in an enhancerless reporter plasmid in H9-derived cardiomyocytes (*** p<0.001 and ** p<0.01 by one-way ANOVA, N=6). **B.** Cardiomyocyte nuclear extract gel shift assay with probes for the reference and AF-risk allele of rs17171731, demonstrating that both probes shift band 1, but only the risk allele shifts band 2. Increasing concentrations of the reference and risk allele unlabeled competitors were added as indicated (10, 25, and 50-fold molar excess). **C.** Gel shift assay of the AF-risk allele using unlabeled competitors (100-fold molar excess) containing one bp changes centered on the rs17171731 SNP, yielding the core sequence element of TGCA for the band 2 gel shift.

**Figure 3.**
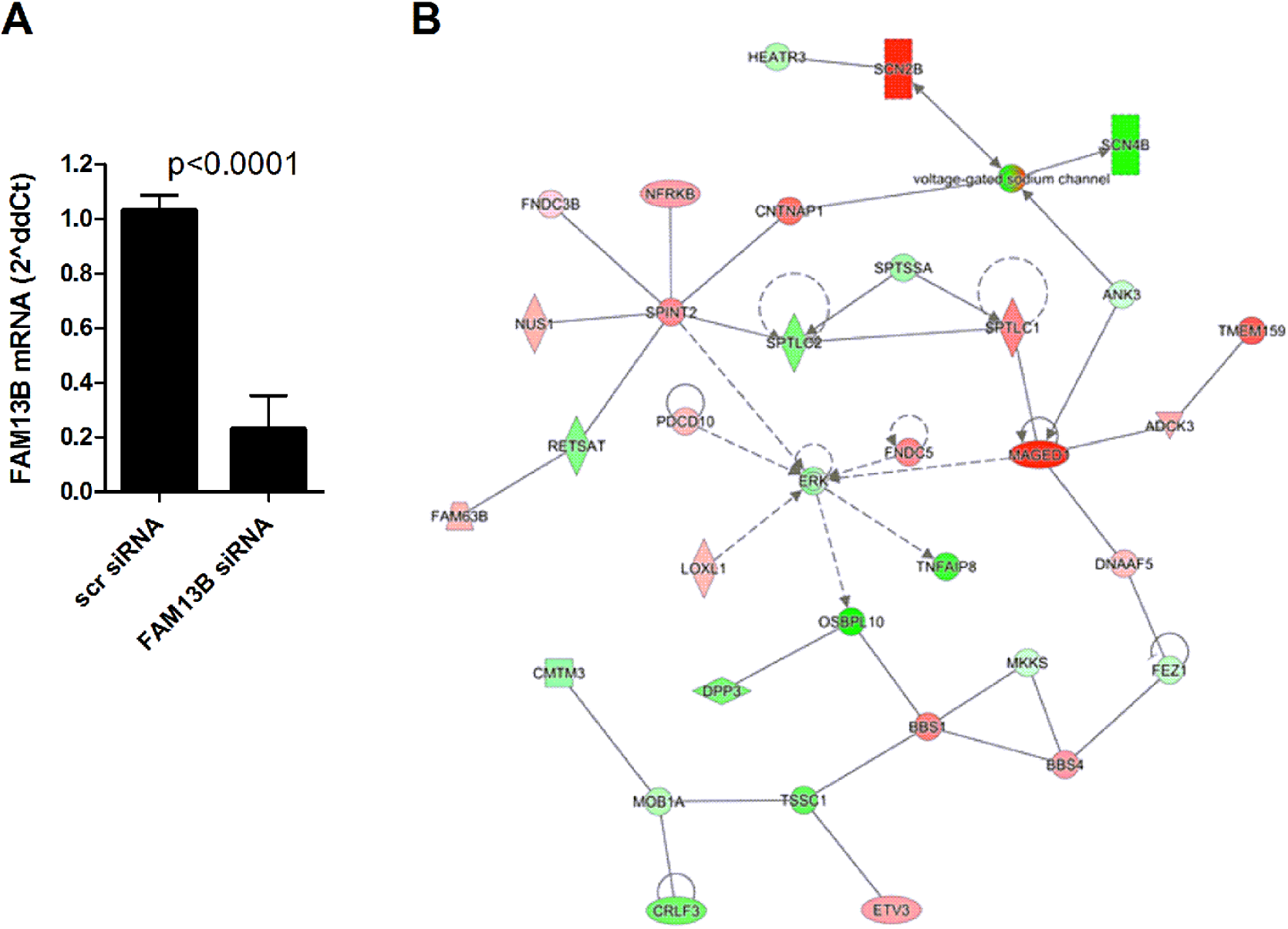
FAM13B Knockdown. **A.** qRT-PCR verification of siRNA *FAM13B* knockdown in iPSC-derived cardiomyocytes normalized to ACTC1 mRNA relative to the scramble siRNA (mean ± SD, p<0.0001 by two-tailed t-test, N=4 for scramble and N=5 for knockdown from 2 independent experiments). **B.** Ingenuity Pathway Analysis of gene expression changes in *FAM13B* knockdown vs. scramble-siRNA control. This was the second strongest empirical network identified (score = 46), and it included the voltage gated sodium channel and the *SCN2B* gene, the 11^th^ strongest regulated gene upon *FAM13B* knockdown. Green indicates repression and red indicates induction upon *FAM13B* knockdown.

**Figure 4.**
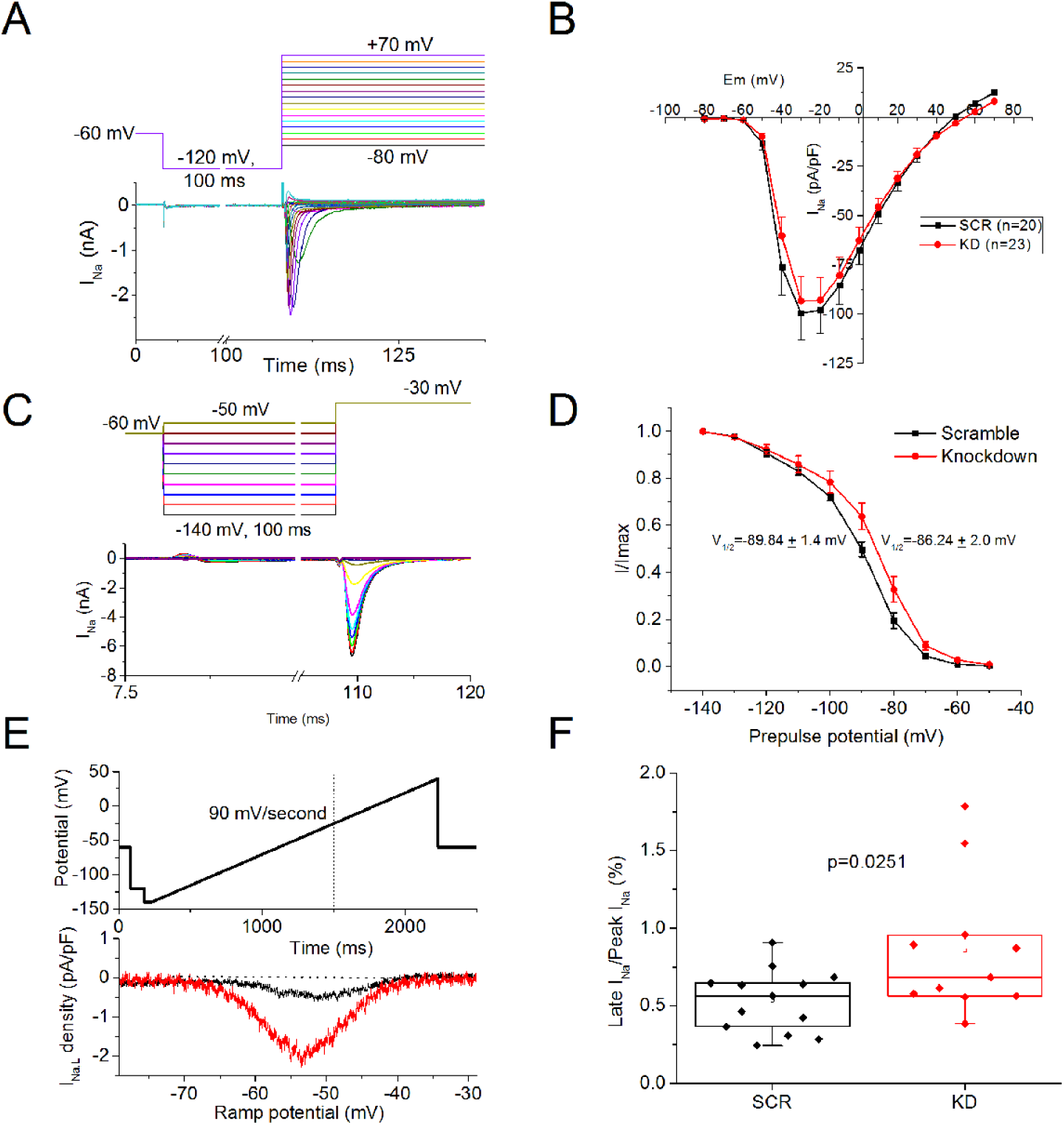
Impact of *FAM13B* knockdown on iPSC-derived cardiomyocyte sodium currents. **A**. Representative I_Na_ traces and voltage-clamp protocol. A 100 ms prepulse to −120 mV was used from a holding potential of −60 mV. Depolarizing voltage steps from −80 to +70 mV were applied in 10 mV increments. **B**. Summary I_Na_-V plot for scramble (n=20) vs. *FAM13B* knockdown (n=23) recordings. **C**. Inactivation protocol and representative I_Na_ traces. As in A, resting potential was −60 mV, with a 100 ms pre-pulse used to partially recover channels from inactivation, using pre-pulse potentials from −140 to −50 mV in 10 mV steps. I_Na_ was recorded with a step to −30 mV. **D**. I_Na_ currents were normalized to those recorded at −140 mV (I/Imax). A Boltzmann function was used to fit the half-inactivation potential (V_½_). A subset of the cells in B were characterized (scramble n=9, knockdown n=7). **E**. Slowly inactivating I_Na_ (I_NaL_) was recorded using a 2s ramp clamp protocol from −140 to +50 mV (90 mV/second). Representative current traces normalized to cell capacitance are shown over the potential range −80 to −30 mV for a scramble (black) and *FAM13B* knockdown (red) cardiomyocyte. The baseline subtracted peak negative amplitude of the current recorded in this range was used as a measure of I_NaL_. **F**. A subset of the cells in B were evaluated using the protocol in **F**. The percent of I_NaL_ to peak I_Na_ is plotted from scramble (n=11) vs. *FAM13B* knockdown (n=13) cells.

### Analysis of *FAM13B-GFP* expression

The FAM13B-EGFP fusion protein expression vector was constructed by VectorBuilder Inc (#VB190426-1100mup). The elongation factor 1α short (EFS) promoter drives the expression of the human FAM13B coding sequence[NM_016603.3] fused at its C-terminus to the eGFP coding sequence, with a Neomycin resistance gene inserted downstream of FAM13B-EGFP sequence connected by T2A Linker.. iCell Cardiomyocytes (Cellular Dynamics) were plated at a density of 5× 10^4^ cells per well into 0.1% Gelatin treated glass-bottom 96-well plate according to the manufacturer’s instruction. The cells were maintained in iCell Cardiomyocytes Maintenance medium. 48 hours later, the cells were transfected with EFS-FAM13B-EGFP plasmid using ViaFect transfection Reagent (Promega Corporation) at a 4:1 reagent-to-DNA ratio (vol:wt) and 0.1 µg plasmid DNA per well. Four to six days later, the cardiomyocytes were fixed with 10% Neutral Buffered Formalin for 10 min at room temperature. After incubation with 0.1% Triton X 100 for 20min to permeabilize the cell membrane, the cells were washed and incubated with Casein Blocker (Thermo Fisher Scientific) for 1 hour at room temperature. The cells were incubated with mouse anti α-actinin primary antibody (1:500 dilution, #A7811, Sigma Aldrich) at 4°C overnight, and then with Alexa Fluor 568 labeled goat anti-mouse IgG secondary antibody (1:500 dilution, #A11004, Thermo Fisher Scientific) for 2 hours at room temperature. Nuclei were counterstained with Hoechst 33342 for 5min and epifluorescent micrographs were obtained.

## Data Access

Variance normalized gene expression levels for each subject are available in the GEO database, accession number GSE69890. LAA eQTL data can be obtained on our web browser: http://afeqtl.lerner.ccf.org/.

## Results

### African descent eQTL analysis for FAM13B gene expression

Our published left atrial appendage RNAseq-eQTL analysis focused on 235 subjects with self-reported European ancestry as well as 30 African American subjects whose ancestry was verified by SNP array principal component analysis, to be admixed or clustered closely with African reference samples. ^4^ In the European ancestry subjects, we showed that the GWAS SNP at the chr. 5q31 locus, rs2040862, was a strong eQTL for expression of the FAM13B gene (p = 7×10^-30^). However, this SNP is in strong linkage disequilibrium (LD) with many other SNPs that have similar eQTL p-values. ^4^ Although our African American cohort was small, we were able to detect a significant eQTL for FAM13B expression. The rs17171731 SNP, located ∼20 kb upstream of the FAM13B transcription start site, stood out with an eQTL p-value ∼10-fold stronger than the other SNPs in this region (Figure 1A, B; p = 2.63×10^-5^). rs17171731 is in high LD with the GWAS SNP in our European ancestry subjects (r^2^ = 0.96). Note, rs17171731 is not included in the 1000 Genomes phase 3 database, but it was present in the phase 1 database used for imputation.

### Functional studies of the FAM13B candidate causal SNP

Based on the cis-eQTL analysis (Figure 1A, B) and DNaseI hypersensitivity in human fetal heart (Figure 1C), we identified rs17171731 as the top candidate causal SNP regulating *FAM13B* gene expression. In order to detect enhancer activity in this region, we subcloned 26-mer oligos containing the reference or risk alleles of rs17171731 (Table 1) into a luciferase reporter plasmid driven by a minimal viral thymidine kinase (TK) promoter (pT81luc). After transfection into cardiomyocytes differentiated from H9 embryonic stem cells, we observed that the risk allele had weak enhancer activity, and the reference allele had 3.3-fold stronger activity vs. the risk allele (p<0.001, Figure 2A).

We then performed electromobility shift assays using ^32^P-labeled 26mer oligos surrounding the rs17171731 SNP (Figure 2B). Incubation of the reference allele probe with human heart nuclear extract yielded one strongly shifted fragment (Band 1), while the risk allele yielded both Band 1 and a smaller shifted fragment (Band 2). Band 1 was effectively competed off of both probes with excess unlabeled reference or risk oligos; however, Band 2 was competed off more effectively with the risk vs. reference oligo. Using a different lot of heart nuclear extract that had more prominent Band 2 vs. Band 1 shifts, along with the risk allele probe, we performed competition studies using nine unlabeled oligos (Table 1) with single bp changes extending ±4 bp from the rs17171731 SNP site (Figure 2C). We determined that the central sequence motif required for the Band 2 shift was TGCA, with the rs17171731 risk allele representing the C of this motif. We searched the JASPAR transcription factor motif database ^11^ for TGCA binding factors with scores >93.5 and narrowed our list to those that were expressed in LA based on our RNAseq data ^4^, which yielded the following eight factors: ATF4, POU2F1, SOX4, ZNF143, SOX8, ABI3, MTF1, and POU2F2. Commercially available antibodies against these proteins did not supershift or block Band 2 formation (data not shown); thus, we did not identify the heart nuclear protein(s) responsible for the Band 2 shift, which presumably mediate the silencing activity of the rs17171731 risk allele.

*FAM13*B (prior gene name C5ORF5 ^5^) is a member of the Rho-GAP gene family. *In silico* analysis of the FAM13B Rho-GAP domain indicates that two highly conserved putative GTPase catalytic residues are altered in the FAM13B protein, potentially yielding a non-catalytic protein. ^5^ To begin to evaluate the mechanism whereby FAM13B impacts atrial arrhythmogenesis, RNAseq and patch clamp studies were performed to assess the effect of *FAM13B* siRNA-transfection vs. a scrambled siRNA on cardiomyocytes derived from iPSCs. Four days after *FAM13B* siRNA transfection, we observed 77% knockdown of *FAM13B* mRNA expression by qPCR (Figure 3A). RNAseq analysis of the *FAM13B* and scrambled siRNA treated cells revealed that expression of 1156 genes was significantly altered after *FAM13B* knockdown (FDR adjusted p<0.1, n=3 per condition, Supplemental Table 1), and the top 20 most significant changes in gene expression are shown in Table 2. While expression levels of prominent atrial ion channel pore subunits were not significantly impacted by *FAM13B* knockdown, expression of *SCN2B*, which encodes a beta-subunit of the cardiac sodium channel, was the 11^th^ strongest regulated transcript, and it was down regulated 2.2-fold by *FAM13B* knockdown (Table 2). Ingenuity Pathway analysis revealed that *SCN2B* was a member of the second strongest empirical network associated with *FAM13B* knockdown (Figure 3B).

**Table 2.**
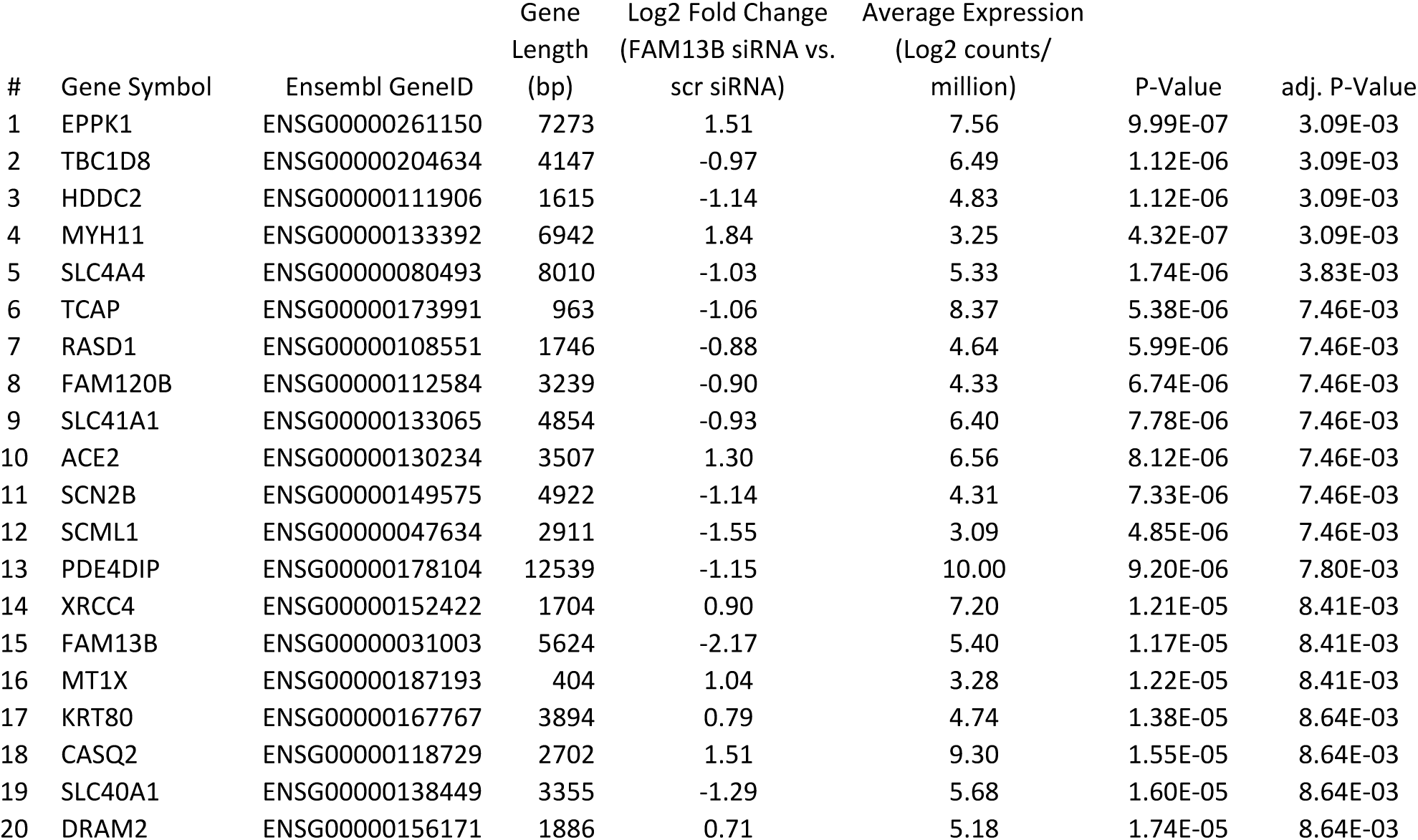
Twenty most significantly regulated genes after FAM13B knockdown.

Patch clamp studies of *FAM13B* knockdown cells focused on measurement of sodium current (I_Na_) density (Figure 4A,B), steady state inactivation (Figure 4C,D) and late sodium current (I_NaL_, Figure 4E,F). *FAM13B* knockdown induced loss of *SCN2B* did not significantly impact the peak sodium current density (Figure 4B). *FAM13B* knockdown increased the V_½_ for the steady state inactivation of the I_Na_ by 2.7 mV (Figure 4D), and although this effect did not reach p<0.05 by t-test, a 2-way ANOVA test showed a significant difference between the curves (p=0.002 for interaction between treatment and voltage). Cell capacitance was not significantly different between groups (KD: 39.5±6.5 pF vs. SCR 36.4±4.5 pF). Late sodium current density (I_NaL_) was evaluated using a ramp clamp protocol (Figure 4E). There was a tendency for I_NaL_ density to be larger in the *FAM13B* knockdown cells compared to the scramble (−1.73±0.33 vs. −1.24±0.13 pA/pF, p=0.20). When subdivided by cell capacitance above and below the median value (37.7 pF), the difference in I_NaL_ density between knockdown and scramble groups was significant for the smaller capacitance cells (p=0.0443), but not for the larger capacitance cells (p=0.879). The ratio of I_NaL_/peak I_Na_ was significantly increased in the *FAM13B* knockdown cells, regardless of cell capacitance (Figure 4F, p=0.025).

To gain more insight into FAM13B mechanism of action, we transfected different cell types with a FAM13B-GFP fusion protein construct. Upon transfection into non-cardiomyocyte HEK293 and human iPS cells, we observed a few GFP+ cells two days after transfection, but these cells were compact, round, and poorly attached, and had died by day four, such that no GFP+ cells remained on the dish. However, after transfection into iCell cardiomyocytes, we observed robust GFP+ staining on the plasma membrane and in sarcomeres (Figure 5). We used antibodies directed to the Z-disc (α-actinin), the sarcoplasmic reticulum (serca2) and we observed the better co-localization with α-actinin (Figure 5).

**Figure 5.**
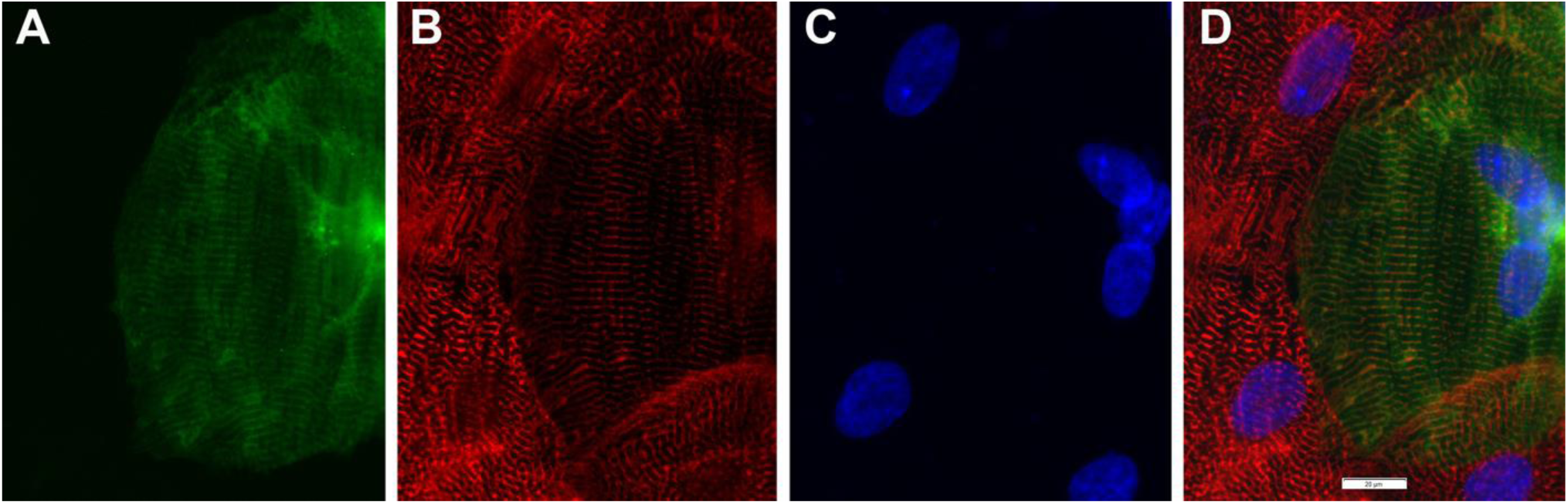
FAM13B-GFP fusion protein expression in iCell cardiomyocytes. **A.** GFP fluorescence of the fusion protein 6 days after transfection, showing expression in a sarcomere pattern and on the plasma membrane. **B.** α-actinin immunofluorescence (red) showing Z-disc localization with the transfected cell surrounded by non-transfected cells. **C.** Hoechst 33342 staining of nuclei. **D.** Merged image, showing FAM13B-GFP expression overlaps with or is adjacent to the Z-disc marker protein α-actinin. 60x objective lens, scale bar show 20 μm.

## Discussion

In our prior LAA RNAseq analysis, the second strongest AF GWAS SNP eQTL was for *FAM13B* at a locus previously attributed to *WNT8A*. ^4^ At this locus, the minor allele of rs2040862 is the AF risk allele, and this allele is associated with decreased expression of *FAM13B*. The *FAM13B* gene encodes an uncharacterized protein containing a Rho GTPase activation protein (RhoGAP) domain. ^5^ The eQTL plots for LAA *FAM13B* expression in European and African American descent subjects and a DNaseI hypersensitivity peak map in fetal heart tissue allowed us to identify rs17171731 as a top candidate functional SNP. Our reporter gene transfection into iPS derived cardiomyocytes show that rs17171731 is a causal variant, with the risk allele having decreased enhancer activity, in the same direction as observed in human LAA tissue. A different SNP at chr. 5q31 (rs1004989) that was previously associated with the electrocardiographic QT-interval was found to be a significant eQTL for *FAM13B* expression in left ventricle tissue samples^12^; however, rs1004989 is not in LD with the best left atrial *FAM13B* eQTL SNP (rs17171731), perhaps illuminating tissue-specific regulatory variants controlling *FAM13B* gene expression.

To begin to assess the functional role of *FAM13B*, we used siRNA knockdown in human cardiomyocytes differentiated from iPSCs. RNAseq analysis of the siRNA knockdown cells showed a striking pattern with extensive co-regulation with more than 1000 other genes, suggesting the FAM13B may be a hub gene that plays a role in regulating LA physiology. Pathway analysis found a strong empirical network that included *SCN2B*, a subunit of the cardiac sodium channel, which we identified as one of the most down-regulated genes. In ventricular cardiomyocytes from normal and heart failure dogs, virally delivered siRNA knockdown of *SCN2B* was associated with increased late sodium current (I_NaL_). ^13^ I_NaL_, resulting from either genetic or environmental influences, has been associated with action potential prolongation in heart failure, and both I_NaL_ ^14^ and mutations in *SCN2B* have been associated with risk of AF. ^15^ Results of our electrophysiology studies in the *FAM13B* knock-down iPSC-derived cardiomyocytes were similar to the effects of *SCN2B* knockdown in canine ventricular cardiomyocytes. ^13^ We noted trends toward a positive shift in the steady state inactivation of the sodium current (Figure 4D) and an increased density of the slowly inactivating sodium current (Figure 4E), resulting in a significantly increased fraction of I_NaL_/peak I_Na_ (Figure 4F). In our study, peak I_Na_ was not different between groups (Figure 4B), but I_NaL_ was increased in smaller (likely atrial and nodal) cells to a greater extent than in the larger capacitance cells. Sodium currents are not identical in atrial and ventricular cardiomyocytes, likely due to differences in sodium channel subunit composition and cellular architecture.

The determinants of peak and late sodium current density are complex, with roles for channel trafficking, CaMKII phosphorylation, subunit interactions, oxidative stress and metabolic status suggested as potential modifiers. We suggest that increased I_NaL_, which indirectly modifies intracellular calcium levels as a result of sodium-calcium exchanger activity, may represent one of the pathways whereby a genetically determined reduction in *FAM13B* abundance increases risk of AF. Other sodium channel (SCN5A) gain of function mutations that increase sodium influx have also been associated with increased risk of cardiac arrhythmia. ^16,17^ Our studies suggest that inhibition of I_NaL_ with new or available drugs, such as ranolazine, may reduce AF risk in patients carrying the *FAM13B* risk allele.

Expression of a FAM13B-GFP fusion protein in iCell cardiomyocytes demonstrated localization both in sarcomeres and on the plasma membrane (Figure 5). The sarcomere staining overlaps or was adjacent to the Z-disc marker protein α-actinin. This pattern may be due to FAM13B located on the Z-disk and/or the T-tubule, which connects the plasma membrane with deep invaginations near the Z-disk. Using the Gene Onloogy list for genes associated with the Z-disk in mice, we found the following other AF GWAS genes ^2^ that share this association: CASQ2, CFL2, FBXO32, KCNN2, MYH7, MYO18B, MYOZ1, SYNE2, SYNPO2L, and TTN

Further understanding of the mechanism by which low expression of FAM13B, along with its binding partners and associated pathways, can predispose to AF susceptibility may illuminate targets for novel therapies. Thus, *FAM13B* is a potential target for drug development to prevent or treat AF.

## Acknowledgements

This work was supported by the National Institutes of Health grant RO1 HL 111314 to MKC, DVW, JB, and JDS, an American Heart Association Strategically Focused Research Network grant 18SFRN34110067 to MKC, DVW, JB and JDS, the NIH National Center for Research Resources for Case Western Reserve University and Cleveland Clinic Clinical and Translational Science Award UL1-RR024989, the Cleveland Clinic Department of Cardiovascular Medicine philanthropy research funds, and the Tomsich Atrial Fibrillation Research Fund. JH was supported by the National Institutes of Health training grant T32 GM 088088. JDS was supported by the Geoffrey Gund Endowed Chair for Cardiovascular Research.

## References

1. Ellinor PT, Yoerger DM, Ruskin JN, MacRae CA. Familial aggregation in lone atrial fibrillation. Hum Genet. 2005;118:179–184.

2. Roselli C, Chaffin MD, Weng L-C, Aeschbacher S, Ahlberg G, Albert CM, Almgren P, Alonso A, Anderson CD, Aragam KG, Arking DE, Barnard J, Bartz TM, Benjamin EJ, Bihlmeyer NA, Bis JC, Bloom HL, Boerwinkle E, Bottinger EB, Brody JA, Calkins H, Campbell A, Cappola TP, Carlquist J, Chasman DI, Chen LY, Chen Y-DI, Choi E-K, Choi SH, Christophersen IE, Chung MK, Cole JW, Conen D, Cook J, Crijns HJ, Cutler MJ, Damrauer SM, Daniels BR, Darbar D, Delgado G, Denny JC, Dichgans M, Dörr M, Dudink EA, Dudley SC, Esa N, Esko T, Eskola M, Fatkin D, Felix SB, Ford I, Franco OH, Geelhoed B, Grewal RP, Gudnason V, Guo X, Gupta N, Gustafsson S, Gutmann R, Hamsten A, Harris TB, Hayward C, Heckbert SR, Hernesniemi J, Hocking LJ, Hofman A, Horimoto ARVR, Huang J, Huang PL, Huffman J, Ingelsson E, Ipek EG, Ito K, Jimenez-Conde J, Johnson R, Jukema JW, Kääb S, Kähönen M, Kamatani Y, Kane JP, Kastrati A, Kathiresan S, Katschnig-Winter P, Kavousi M, Kessler T, Kietselaer BL, Kirchhof P, Kleber ME, Knight S, Krieger JE, Kubo M, Launer LJ, Laurikka J, Lehtimäki T, Leineweber K, Lemaitre RN, Li M, Lim HE, et al. Multi-ethnic genome-wide association study for atrial fibrillation. Nat Genet. 2018;50:1225–1233.

3. Nielsen JB, Thorolfsdottir RB, Fritsche LG, Zhou W, Skov MW, Graham SE, Herron TJ, McCarthy S, Schmidt EM, Sveinbjornsson G, Surakka I, Mathis MR, Yamazaki M, Crawford RD, Gabrielsen ME, Skogholt AH, Holmen OL, Lin M, Wolford BN, Dey R, Dalen H, Sulem P, Chung JH, Backman JD, Arnar DO, Thorsteinsdottir U, Baras A, O’Dushlaine C, Holst AG, Wen X, Hornsby W, Dewey FE, Boehnke M, Kheterpal S, Mukherjee B, Lee S, Kang HM, Holm H, Kitzman J, Shavit JA, Jalife J, Brummett CM, Teslovich TM, Carey DJ, Gudbjartsson DF, Stefansson K, Abecasis GR, Hveem K, Willer CJ. Biobank-driven genomic discovery yields new insight into atrial fibrillation biology. Nat Genet. 2018;1234–1239.

4. Hsu J, Gore-panter S, Tchou G, Castel L, Lovano B, Moravec CS, Pettersson GB, Eric E, Gillinov AM, Mccurry KR, Smedira NG, Barnard J, Wagoner DR Van, Chung MK. Genetic Control of Left Atrial Gene Expression Yields Insights into the Genetic Susceptibility for Atrial Fibrillation. Circ Genom Precis Med. 2018;11:e00210.

5. Lai F, Godley LA, Fernald AA, Orelli BJ, Pamintuan L, Zhao N, Le Beau MM. cDNA cloning and genomic structure of three genes localized to human chromosome band 5q31 encoding potential nuclear proteins. Genomics. 2000;70:123–30.

6. Cohen M, Reichenstein M, der Wind A, Heon-Lee J, Shani M, Lewin HA, Weller JI, Ron M, Seroussi E. Cloning and characterization of FAM13A1—a gene near a milk protein QTL on BTA6: evidence for population-wide linkage disequilibrium in Israeli Holsteins. Genomics. 2004;84:374–383.

7. Marchini J, Howie B, Myers S, McVean G, Donnelly P. A new multipoint method for genome-wide association studies by imputation of genotypes. Nat Genet. 2007;39:906–13.

8. Shabalin A a. Matrix eQTL: ultra fast eQTL analysis via large matrix operations. Bioinformatics. 2012;28:1353–8.

9. Storey JD, Tibshirani R. Statistical significance for genomewide studies. Proc Natl Acad Sci. 2003;100:9940–9945.

10. Nordeen SK. Luciferase reporter gene vectors for analysis of promoters and enhancers. Biotechniques. 1988;6:454–458.

11. Mathelier A, Fornes O, Arenillas DJ, Chen C, Denay G, Lee J, Shi W, Shyr C, Tan G, Worsley-Hunt R, others. JASPAR 2016: a major expansion and update of the open-access database of transcription factor binding profiles. Nucleic Acids Res. 2016;44:D110–D115.

12. Arking DE, Pulit SL, Crotti L, der Harst P, Munroe PB, Koopmann TT, Sotoodehnia N, Rossin EJ, Morley M, Wang X, others. Genetic association study of QT interval highlights role for calcium signaling pathways in myocardial repolarization. Nat Genet. 2014;46:826–836.

13. Mishra S, Undrovinas NA, Maltsev VA, Reznikov V, Sabbah HN, Undrovinas A. Post-transcriptional silencing of SCN1B and SCN2B genes modulates late sodium current in cardiac myocytes from normal dogs and dogs with chronic heart failure. Am J Physiol Circ Physiol. 2011;301:H1596--H1605.

14. Fischer TH, Herting J, Mason FE, Hartmann N, Watanabe S, Nikolaev VO, Sprenger JU, Fan P, Yiao L, Popov A-F, others. Late INa increases diastolic SR-Ca2+-leak in atrial myocardium by activating PKA and CaMKII. Cardiovasc Res. 2015;107:184–196.

15. Watanabe H, Darbar D, Kaiser DW, Jiramongkolchai K, Chopra S, Donahue BS, Kannankeril PJ, Roden DM. Mutations in sodium channel $β$1-and $β$2-subunits associated with atrial fibrillation. Circ Arrhythmia Electrophysiol. 2009;2:268–275.

16. Tian X-L, Yong SL, Wan X, Wu L, Chung MK, Tchou PJ, Rosenbaum DS, Van Wagoner DR, Kirsch GE, Wang Q. Mechanisms by which SCN5A mutation N1325S causes cardiac arrhythmias and sudden death in vivo. Cardiovasc Res. 2004;61:256–267.

17. Blana A, Kaese S, Fortmüller L, Laakmann S, Damke D, van Bragt K, Eckstein J, Piccini I, Kirchhefer U, Nattel S, others. Knock-in gain-of-function sodium channel mutation prolongs atrial action potentials and alters atrial vulnerability. Hear Rhythm. 2010;7:1862–1869.

